# Host glutathione is required for *Rickettsia parkeri* to properly septate, avoid ubiquitylation, and survive in macrophages

**DOI:** 10.1101/2023.10.02.560592

**Authors:** Han Sun, Anh Phuong Luu, Meggie Danielson, Thaomy T. Vo, Thomas P. Burke

## Abstract

Spotted fever group *Rickettsia* obligately reside in the cytosol where they parasitize over fifty metabolites from their hosts. However, the role for metabolite acquisition in pathogenesis remains unclear. Here, we find that depletion of the abundant low molecular weight thiol glutathione led to an impaired ability of *Rickettsia parkeri* to form plaques. Super-resolution microscopy revealed that glutathione depletion with buthionine sulfoximine (BSO) in endothelial or epithelial cells led to the formation of bacterial chains that increased in length over time. Chained bacteria had fewer actin-tails and were impeded for their ability to spread from cell to cell. Glutathione depletion also caused an increased frequency of colocalization between the bacterial surface and polyubiquitin. *R. parkeri* was significantly more restricted upon glutathione depletion in primary macrophages than in epithelial cells in a mechanism that avoided activating the inflammasome and the production of type I interferon. Together, these data suggest that host glutathione is critical for rickettsial septation, actin-based motility, avoiding ubiquitylation, and survival in immune cells.

## Introduction

Spotted fever group (SFG) *Rickettsia* species are dangerous agents of disease in the United States and worldwide (1). SFG *R. rickettsii* is the causative agent of Rocky Mountain spotted fever (RMSF), a disease characterized by fever of over 104° F, maculopapular rash, severe headache, and in some cases death (2). Case-fatality rates for RMSF were as high as 65-80% prior to antibiotics (3) and can be fatal even after antibiotic treatment (4). The SFG pathogen *R. parkeri* is the causative agent of mild spotted fever disease in North and South America (5–7) and has a highly similar genome to *R. parkeri*, with some virulence genes containing >99% sequence alignment, with the major difference being a single 33.5 kb locus that is absent in *R. rickettsii* (5). *R. parkeri* rickettsiosis is characterized by a skin lesion (eschar) at the site of tick bite, fever of 103° F, rash, and headache (6, 7). Spotted fever has increased in incidence from approximately 500 cases in the year 2000 to 6,248 cases in 2017, and disease occurs in every state of the continental United States (1). Spotted fever and other tick-borne diseases are increasing in prevalence as the range of ticks expands, which may be fueled by climate change (8). The increasing incidence combined with difficulties in recognizing spotted fever clinically (9) illustrate a critical need to better understand molecular mechanisms of disease caused by SFG rickettsial pathogens.

*Rickettsia* and their close relatives are obligate to the host cell cytosol, where they parasitize over 50 nutrients (10, 11). *Rickettsia* have highly reduced genomes with degraded biosynthetic pathways, including loss of enzymes required for glycolysis and the tricarboxylic acid cycle, as well as biosynthesis of fatty acids, isoprenoids, vitamins, cofactors and amino acids, and the abundant low molecular weight thiol glutathione (5, 10, 12). Glutathione prevents oxidative damage by reducing reactive oxygen species (ROS), and it can also function as a cysteine source after conversion to methionine. S-glutathionylation of cysteine can function as a post-translational modification that alters protein activity. *Rickettsia* encode a conserved glutathione S-transferase, an enzyme that catalyzes the addition of glutathione to substrates (10, 12), however the role for glutathione in intracellular growth and pathogenesis remains unclear.

Glutathione is critical for pathogenesis of other phylogenetically distinct cytosol-dwelling pathogens, including *Listeria monocytogenes, Burkholderia pseudomallei* and *Francisella* species. An extracellular histidine kinase domain of *B. pseudomallei* binds glutathione initiate expression of type VI secretion system (13). Inhibition of glutathione synthesis by buthionine sulfoximine (BSO), which is a potent, specific, non-toxic, and irreversible inhibitor of y-glutamylcysteine synthetase, blocks *B. pseudomallei* virulence gene expression and inhibits cell to cell spread (13). In the case of *Listeria monocytogenes*, the bacteria sense host methylgloxyl, resulting in the bacterial production of glutathione that activates the master virulence gene regulator PrfA (14–16). In *Francisella* species, glutathione is imported and degraded as a sulfur source (17–19). Other pathogens, including *Streptococcus* species and *Hemophilus influenzae* co-opt host glutathione to defend against oxidative stress (20, 21). These studies support an emerging concept that glutathione plays important yet distinct roles in virulence of intracellular bacteria.

Here, we investigated the role for glutathione in the intracellular growth and survival of an obligate cytosolic pathogen, *R. parkeri*. We find that it is required for specific aspects of intracellular growth in epithelial and endothelial cells, including proper septation, actin-based motility, and avoiding ubiquitylation. We also observe that glutathione is significantly more important for *R. parkeri* survival in macrophages than in epithelial or endothelial cells. These findings establish a critical role for host glutathione in *Rickettsia* pathogenesis and contribute to an emerging paradigm that cytosol-dwelling bacterial pathogens sense the host reducing environment for specific aspects of virulence.

## Results

### Glutathione is required for proper plaque formation and septation of *R. parkeri* in epithelial and endothelial cells

To investigate the role for glutathione in *R. parkeri* pathogenesis, we depleted glutathione with BSO in Vero cells, which are primate epithelial cells that are highly permissive to infection and measured *R. parkeri* plaque formation at 7 days postinfection (dpi). To discriminate between any potential effects that BSO had on the bacteria versus on the host cells, BSO was added to Vero cells overnight, then washed from the cells 30 mins prior to infection with *R. parkeri*. At 7 dpi, infected cells were fixed and stained with crystal violet. High multiplicities of infection (MOIs) caused host cell lysis, yet *R. parkeri* was >500-fold attenuated in its ability to form plaques at lower MOIs in BSO-treated cells (**Fig. 1A**). To determine if the anti-rickettsial effect was due to BSO acting on the host cells or potentially on the bacteria, *R. parkeri* was incubated with BSO prior to infection of untreated Vero cells. Incubation of *R. parkeri* with BSO did not affect PFUs as compared to untreated controls (**Fig. 1A**), suggesting that BSO was acting on the host cells and not on *R. parkeri* directly. These data suggest that *R. parkeri* requires host glutathione for plaque formation.

**Fig. 1:**
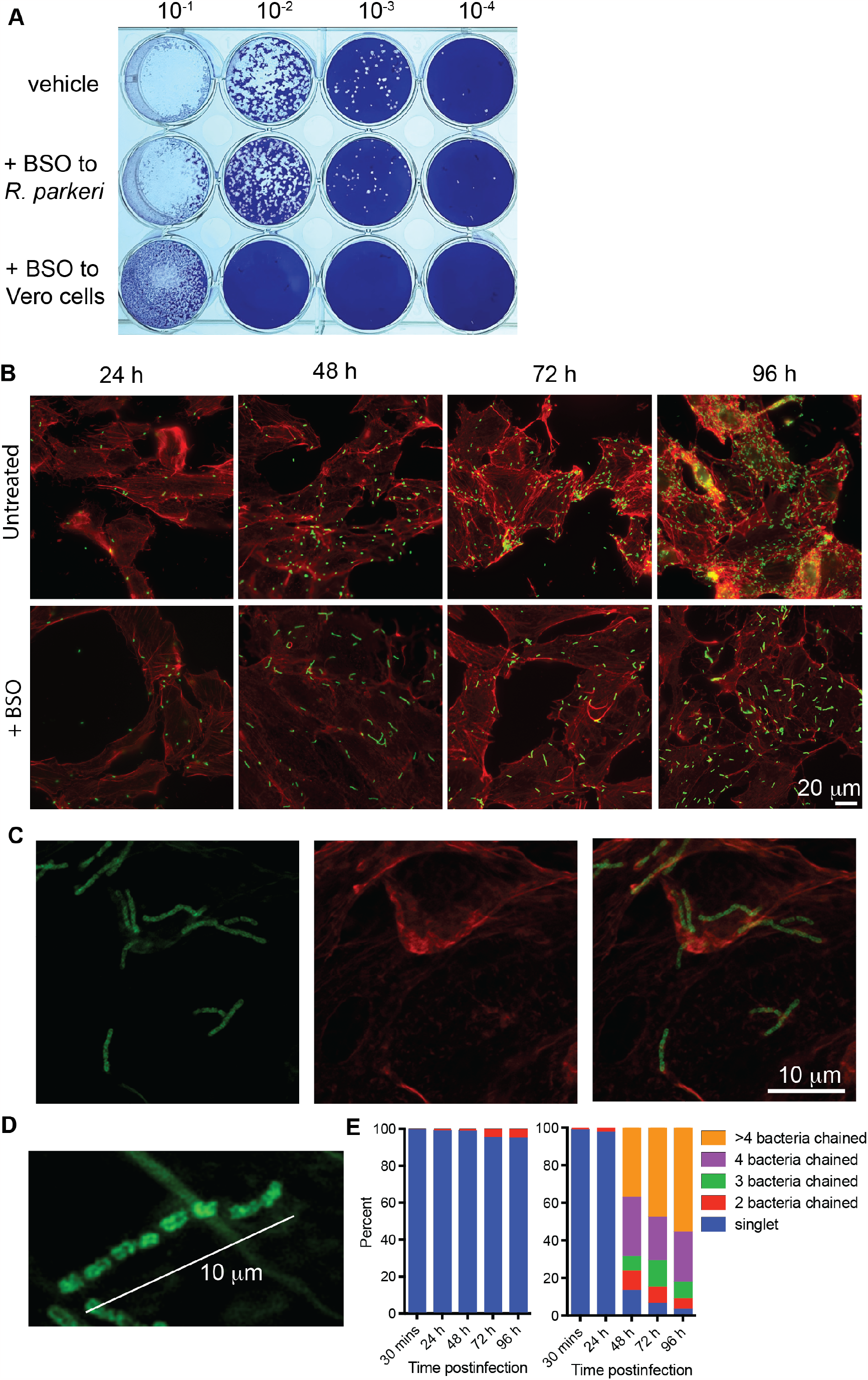
Glutathione is required for proper plaque formation and septation of *R. parkeri* in endothelial and epithelial cells. **A**) Vero cells were infected with *R. parkeri* at the indicated dilutions. For treatment of Vero cells, 2 mM BSO was added 16 h prior to infection, removed during infection, and added 30 again at mpi. For treatment of *R. parkeri*, bacteria were treated on ice for 15 m prior to infection with 2 mM BSO. Avicel was added at 24 hpi and cells were fixed at 7 dpi for crystal violet staining. Image is representative of three independent experiments. **B**) Representative immunofluorescence microscopy images of Vero cells infected with *R. parkeri* on coverslips in 24 well plates. **C**,**D)** Representative images of 63x confocal super-resolution microscopy of Vero cells treated with 2 mM BSO and infected with *R. parkeri*, at 48 hpi. For all panels, cells were infected at an MOI of 1 and fixed with 4% paraformaldehyde at the indicated times. Coverslips were then stained with phalloidin (red) and with an anti-*Rickettsia* antibody (green). **E)** Quantification of images from HMEC-1 cells. Data are representative of at least three separate experiments.

To better understand the mechanism by which glutathione is required for plaque formation, we analyzed infection with immunofluorescence microscopy. In Vero cells treated with BSO, *R. parkeri* had normal morphology early during infection (<24 h), but the bacteria had altered morphology as time increased. Doublet bacteria were significantly more prevalent at 24 hpi in BSO-treated cells, and from 48-96 hpi the bacteria were observed in chains of 3, 4, and >4 bacteria (**Fig. 1B**). To better discern the morphology of *R. parkeri* in BSO-treated cells we next employed super-resolution microscopy. Airyscan confocal laser scanning microscopy revealed that many bacterial chains measured over 10 µM in length (**Fig. 1C**). Moreover, distinct separation was observed within the chained bacteria, suggesting that the chaining was due to a defect in septation. In contrast, bacteria in untreated cells were >90% single rods throughout the infection that consistently measured ~1 µM in length (**Fig. 1E**). This suggests that host glutathione is required for proper septation of *R. parkeri*.

### Glutathione is required for proper actin-based motility of *R. parkeri*

It remained unclear if host glutathione was required for actin-based motility, which is critical for cell-cell spread and contributes to plaque formation. *R. parkeri* undergoes two temporally distinct phases of actin-based motility, an early phase mediated by RickA that results in short, curly actin tails, and a later phase mediated by Sca2 that results in long tails and is the major mediator of cell-cell spread (22–25). We quantified the frequency of actin-tail formation and actin colocalization in untreated and glutathione depleted cells. At 30 mpi, BSO did not affect the frequency of absent actin, polar actin, diffuse actin, curly tails, or straight tails as compared to untreated cells. At 48 hpi, chained bacteria in BSO-treated cells had observable actin tails but with reduced frequency as compared to untreated (**Fig. 2C**). Polar actin was often observable within chains of bacteria, suggesting that these bacteria had defects in septation and were not filamentous with a shared cytosol (**Fig. 2B**). At 72 hpi, *R. parkeri* in BSO-treated cells were largely absent of actin, as compared to untreated cells (**Fig. 2 C**). Lastly, as many *Rickettsia* species target endothelial cells during systemic infection, we determined the effects of BSO on infected human microvascular endothelial cells (HMEC-1). Similar results were observed between in HMEC-1 cells as Vero cells, including defects in septation and actin-based motility (**Fig. 2D**). These data demonstrate that host glutathione is required for proper actin-based motility of *R. parkeri* and suggest that defects in cell-to-cell spread in BSO-treated cells may contribute to the inability of *R. parkeri* to form plaques.

**Fig. 2:**
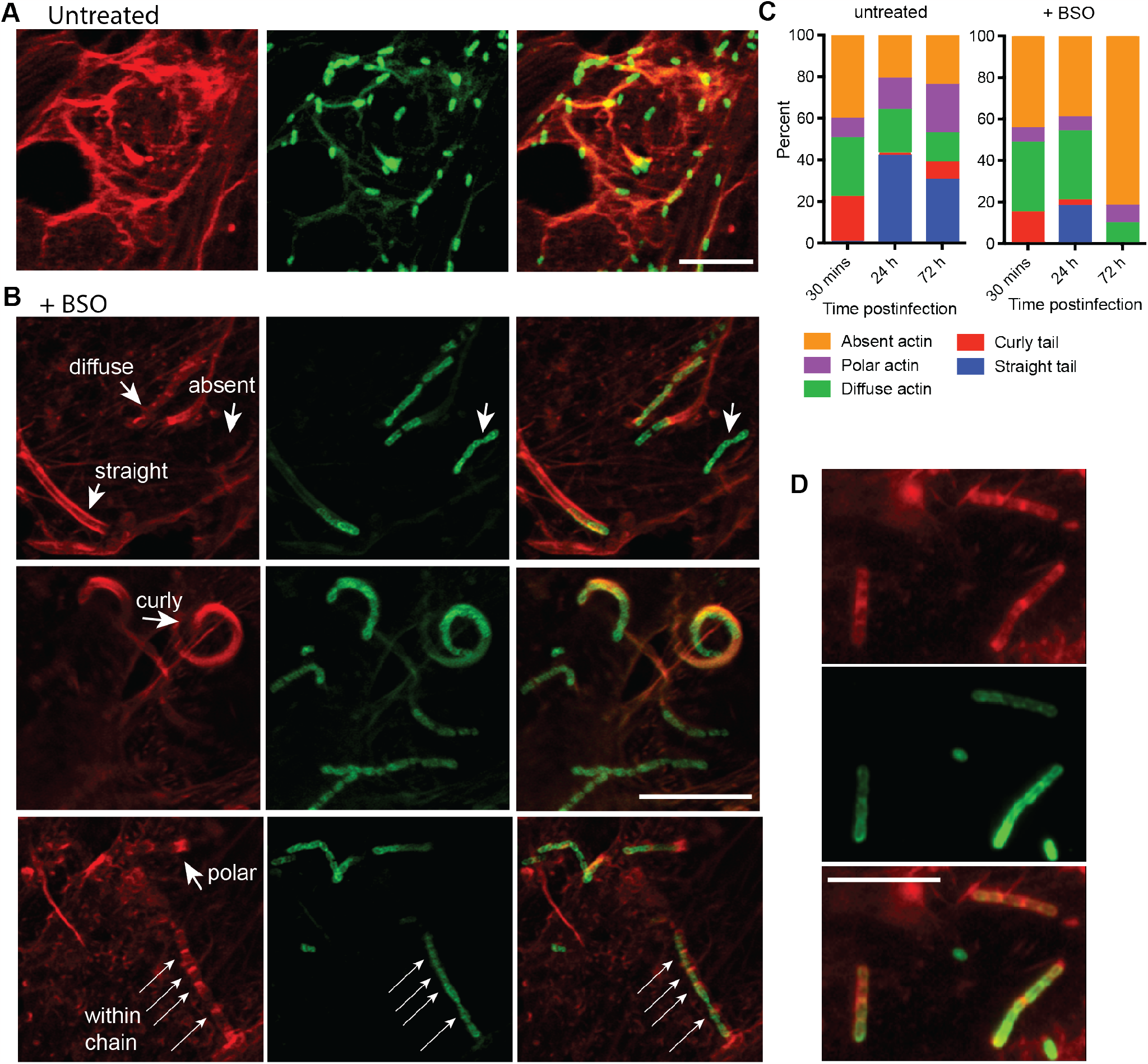
Host glutathione is required for proper actin-based motility of *R. parkeri*. **A**) Representative images using 20x confocal immunofluorescence microscopy of Vero cells infected with *R. parkeri* at an MOI of 1 and imaged at 48 hpi. Scale bar = 10 µm. **B**) Representative images using 63x confocal immunofluorescence microscopy of Vero cells infected with *R. parkeri* at an MOI of 1 and imaged at 48 hpi. 2 mM BSO was added prior to infection overnight. Scale bar = 10 µm. **C)** Quantification of actin colocalization around *R. parkeri*. Each time point is an average of at least 4 separate experiments totaling >1,000 bacteria. **D)** Representative images of HMEC-1 cells treated with 2 mM BSO overnight and infected with *R. parkeri*, with 63x confocal microscopy at 48 hpi. For all microscopy experiments, coverslips containing infected cells were fixed with 4% PFA and stained with phalloidin (red), anti-*Rickettsia* (green). Data represent 3 independent experiments.

**Glutathione depletion restricts *R. parkeri* in macrophages but does not cause significantly enhanced inflammasome activation or interferon production**.

Many spotted fever group *Rickettsia* associate with leukocytes throughout their infectious lifecycles, including macrophages and neutrophils in the skin and in internal organs upon dissemination (7, 26–29). We therefore sought to determine if glutathione was required for *R. parkeri* survival in primary macrophages. In bone marrow-derived macrophages (BMDMs), depletion of glutathione with BSO led to a reduction in recoverable PFUs that was more severe than in epithelial cells (**Fig. 3A**). The loss of recoverable PFUs could be due to either bacterial restriction or alternatively due to an inability of surviving bacteria to form plaques in the PFU assay. To discriminate between these possibilities, we analyzed infected BMDMs with immunofluorescence microscopy. This revealed that *R. parkeri* was largely absent from BMDMs at 48 hpi (**Fig. 3B**), suggesting that glutathione was essential for survival in macrophages.

**Fig. 3:**
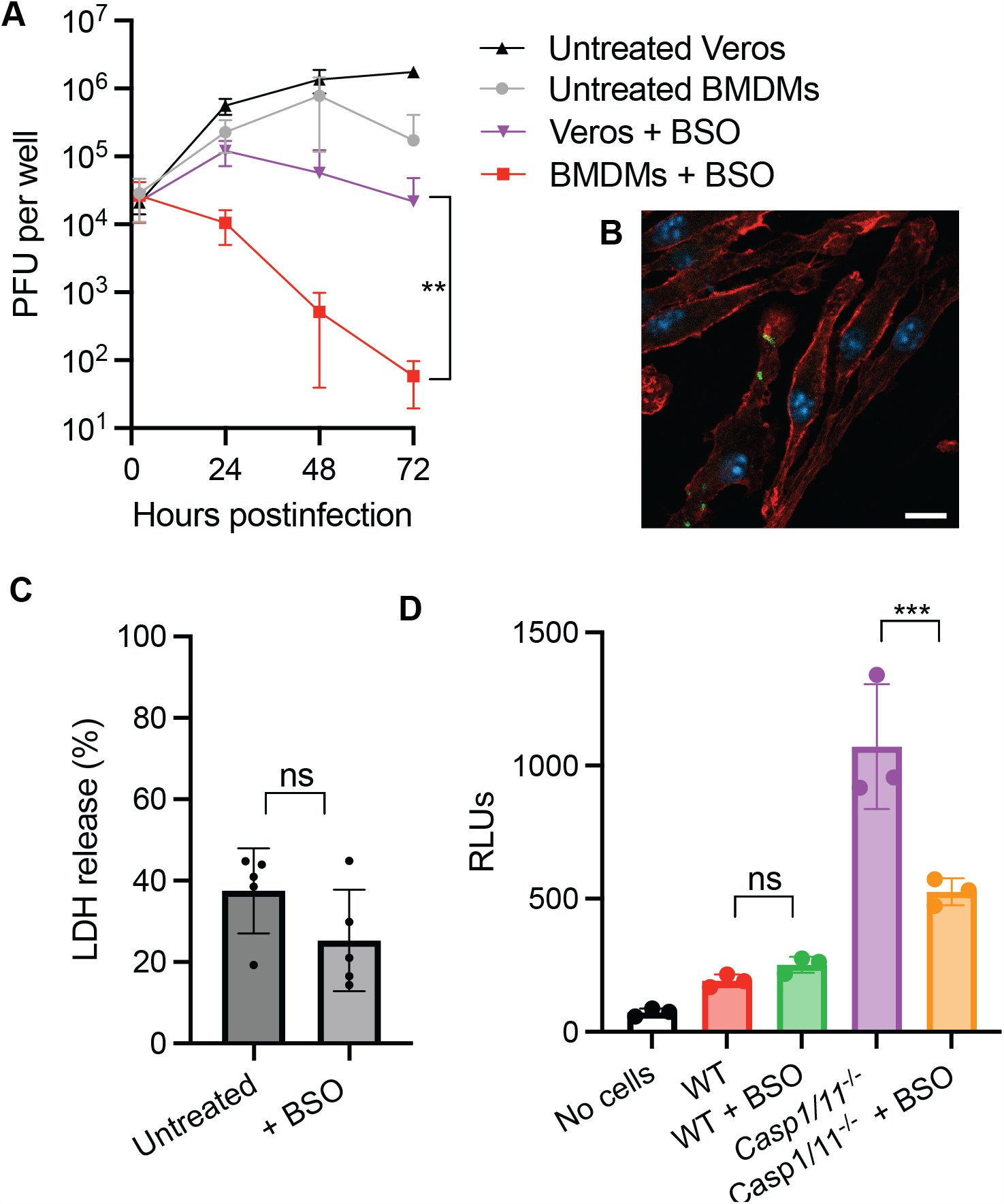
Host glutathione is required for *R. parkeri* survival in macrophages via a non-bacteriolytic restriction mechanism. **A**) *R. parkeri* abundance WT murine BMDMs or Vero cells infected at an MOI of 1. Each data point is the average of 3 independent experiments. Bars denote SD. Statistics used a two-way ANOVA. **B**) Representative images using 60x confocal microscopy of BMDMs treated with 2 mM BSO overnight, infected with R. parkeri at an MOI of 1 and imaged at 48 hpi. **C**) Quantification of host cell death during *R. parkeri* infection of WT BMDMs. LDH release was measured at 24 hpi upon *R. parkeri* infection at an MOI of 1. Data are the compilation of three separate experiments and are expressed as means + SD. Statistical analyses in used a two-tailed Student’s T test. **D**) Measurement of IFN-I in supernatants of BMDMs infected with *R. parkeri* at an MOI of 1. Supernatants were harvested at 24 hpi and used to stimulate a luciferase-expressing cell line (ISRE) and relative light units (RLU) were measured and compared between treated and untreated cells. Statistical comparisons were made use a Student’s two-way T-Test. Data are the compilation of three separate experiments and are expressed as means + SD. **p<0.01, ns=not significant.

It remained unclear if restriction in macrophages was the result of bacteriolysis. We previously described how lysis of *R. parkeri* in wild type (WT) macrophages by guanylate binding proteins (GBPs) leads to caspase 11 inflammasome activation and pyroptosis (30). However, in caspase 1 and 11 deficient BMDMs (*Casp1/11*^-/-^), killing of *R. parkeri* by GBPs instead releases DNA that activates the DNA sensor cGAS, causing release of type I interferon (IFN-I, (30)). To determine if glutathione is required for protecting against bacteriolysis, we measured host cell death in WT macrophages and IFN-I release in *Casp1/11*^-/-^ macrophages in the presence or absence of BSO. In WT macrophages, BSO treatment resulted in similar LDH release than in untreated cells (**Fig. 3C**), suggesting that glutathione was not required for protecting against bacteriolysis. Infection of BSO-treated and untreated BMDMs resulted in similar amounts of IFN-I production in WT cells, while the amount of IFN-I production was reduced in BSO-treated *Casp1/11*^-/-^ cells as compared to untreated cells (**Fig. 3D**). The explanation for reduced IFN-I in BSO-treated *Casp1/11*^*-/-*^ is unclear, however this could be related to a stronger restriction of the bacteria, which we previously reported. Nevertheless, the lower IFN-I upon BSO-treatment suggests that glutathione depletion is not eliciting higher bacteriolysis and cGAS activation. Together, these data demonstrate that restriction of *R. parkeri* in macrophages is via a non-lytic mechanism that does not activate cytosolic pattern recognition receptors including caspase 11 or cGAS.

### Host glutathione is required for *R. parkeri* to avoid targeting by ubiquitin

*R. parkeri* mutants that are unable to avoid targeting by ubiquitin are restricted in BMDMs but not in Vero cells (31, 32). Since restriction of *R. parkeri* by BSO in macrophages did not result in bacteriolysis, we hypothesized that restriction may involve ubiquitin-mediated restriction of the bacteria. We therefore examined whether glutathione depletion increased ubiquitin recruitment to *R. parkeri* in Vero cells and BMDMs. Immunofluorescence microscopy revealed that ~12% of the bacteria in BSO-treated Vero cells were positive for polyubiquitin, which was significantly more than the ~1% observed in untreated cells (**Fig. 4A**). These data demonstrate that host glutathione is required for *R. parkeri* avoidance of ubiquitylation and are aligned with our findings that restriction is via a non-bacteriolytic mechanism.

**Fig. 4:**
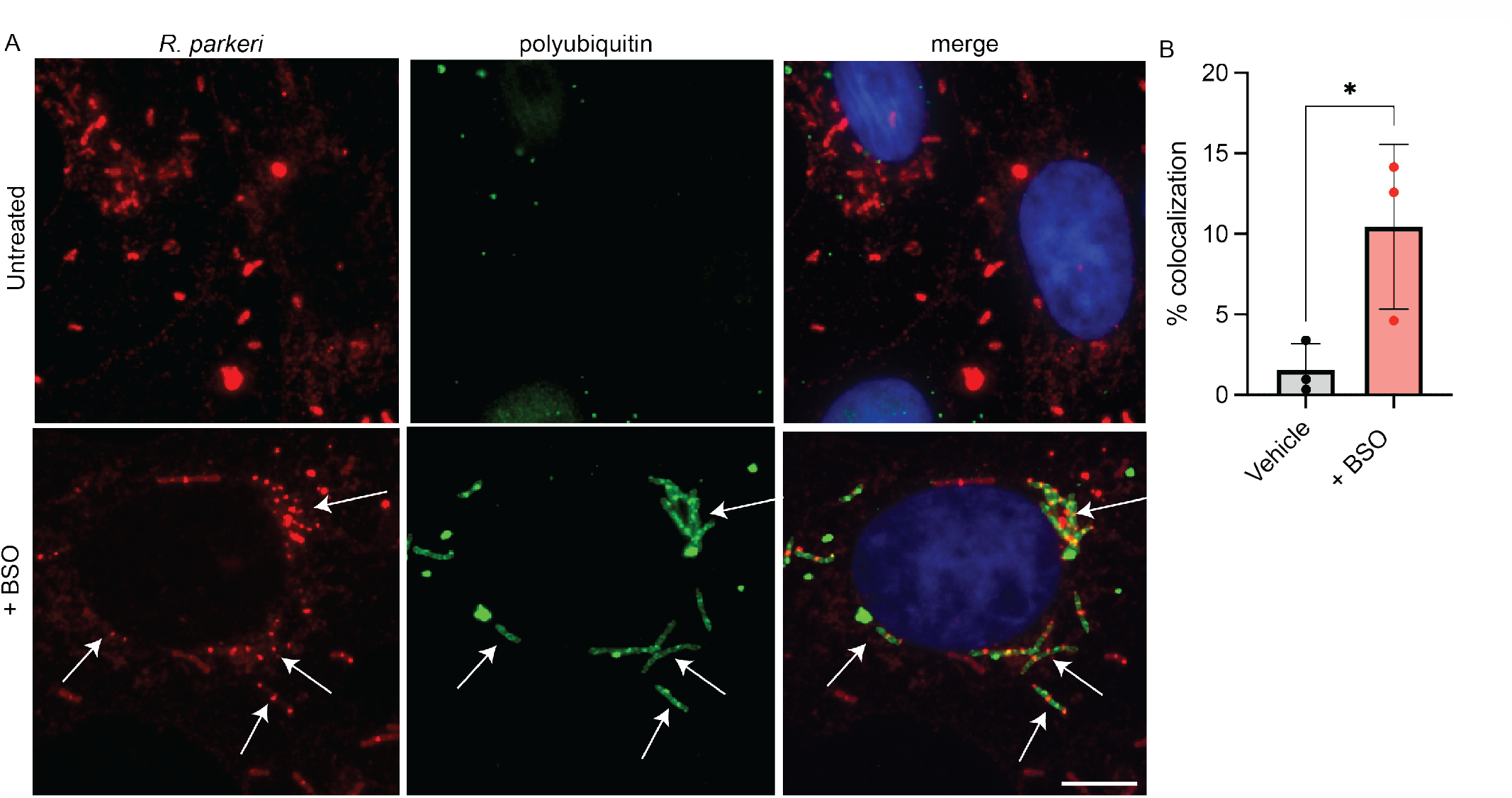
Host glutathione is required for *R. parkeri* to avoid ubiquitylation. **A**) Representative images of ubiquitin colocalization using immunofluorescence microscopy of Vero cells infected with *R. parkeri* at an MOI of 1 and fixed at 48 hpi. Cells treated with 2 mM BSO were incubated overnight with the drug. Cells were stained with FK1 anti-polyubiquitin (green), anti-*Rickettsia* I7 (red); and DAPI (blue). Scale bar = 10 μm. **B**) Quantification of ubiquitin colocalization at 48 hpi in Vero cells. Each data point is an independent experiment with at least 4 images and each image contained at least 50 bacteria. Data are expressed as means + SD. Statistical analysis used a two-tailed Student’s T test; *p<0.05.

## Discussion

Parasitizing metabolites from hosts is a critical aspect of pathogenesis for intracellular bacteria. As obligate intracellular pathogens, the *Rickettsia* steal over fifty host nutrients from their hosts and evolved a reduced genome size that has lost the ability to synthesize many metabolites *de novo*, including glutathione (10). In this study we find that host glutathione is required for *R. parkeri* septation, actin-based motility, avoiding ubiquitylation, and survival in macrophages. This reveals that host glutathione plays a critical role in specific aspects of *R. parkeri* pathogenesis and add to a growing paradigm that the reducing environment of the cytosol is a virulence cue for bacteria that reside in this compartment.

Glutathione is an abundant reducing agent of the host cytosol that can detoxify reactive oxygen species, alter proteins post-translationally, and be used as a cysteine source. Glutathione is used as a cue for *Burkholderia* to initiate virulence, whereby the extracellular domain of the histidine kinase VirA is reduced by glutathione, breaking it from a dimer to a monomer and resulting in transcription of the type VI secretion system (13). Additionally, *Listeria* senses the host reducing environment, synthesizes its own glutathione that post-translationally activates the master virulence transcription factor PrfA (14, 15). Our findings reveal that host glutathione contributes *R. parkeri* pathogenesis, which could be mediated through either direct post-translational modification of factors involved in septation, or through virulence gene expression.

These observations across multiple phylogenetically distinct species suggests a paradigm that glutathione is a host-associated molecular pattern (HAMP) that is sensed by cytosolic microbes for virulence. We propose that the concept of HAMPs is similar to but on the opposite side of the coin as PAMPS (pathogen-associated molecular patterns). PAMPs are conserved molecular features of pathogens, such as flagellin, lipopolysaccharide, lipoproteins, and cyclic dinucleotides that are sensed by host cells as a signature of infection. Detection of PAMPs by host pattern recognition receptors (PRRs) leads to a host defense response such as upregulation of antimicrobial genes and programmed cell death. PAMPs and PRRs are well characterized; however, HAMPs are less well understood. Pathogens that reside directly in the cytosol such as *Rickettsia, Listeria, Burkholderia, Shigella* species, and *Mycobacterium marinum*, are ideal systems to study HAMPs because they initiate actin-based motility and other virulence genes only upon cytosolic entry. Identifying other HAMPs will increase our understanding of bacterial pathogenesis and potentially lead to new strategies for therapeutic targeting.

## Methods

### Preparation of *R. parkeri*

*R. parkeri* strain Portsmouth was originally obtained from Christopher Paddock (Centers for Disease Control and Prevention). Bacteria were amplified by infecting confluent T175 flasks of female African green monkey kidney epithelial Vero cells were obtained from the UC Berkeley Cell Culture Facility, where they were tested for mycoplasma contamination, and were authenticated by mass spectrometry experiments. Vero cells were grown in DMEM (Gibco 11965-092) with glucose (4.5 g/L) supplemented with 2% fetal bovine serum (FBS, GemCell) with 5 x 10^6^ *R. parkeri* per flask. Infected cells were scraped and collected at 5 dpi when ~90% of cells were highly infected. Scraped cells were centrifuged at 12,000 x G for 20 min at 4°C. Pelleted cells were then resuspended in K-36 buffer (0.05 M KH_2_PO_4_, 0.05 M K_2_HPO_4_, 100 mM KCl, 15 mM NaCl, pH 7) and dounced (60 strokes) at 4°C. The solution was then centrifuged at 200 x G for 5 min at 4°C to pellet host cell debris. Supernatant containing *R. parkeri* was overlaid on a 30% MD-76R (Merry X-Ray) solution. Gradients were centrifuged at 18,000 rpm in an SW-28 ultracentrifuge swinging bucket rotor (Beckman/Coulter) for 20 min at 4°C to separate host cells debris. Bacterial pellets were resuspended in brain heart infusion (BHI) media (BD, 237500) and stored at -80°C.

Titers were determined via plaque assays by serially diluting the bacteria in 6-well plates containing confluent Vero cells. Plates were then spun for 5 min at 300 x G in an Eppendorf 5810R centrifuge. At 24 hpi, the media from each well was aspirated and the wells were overlaid with 4 ml/well DMEM with 5% FBS and 0.7% agarose (Invitrogen, 16500-500). At 7 dpi, cells were fixed, stained, and counted as described below.

### Deriving bone marrow macrophages

To obtain bone marrow, male or female mice were euthanized, and femurs, tibias, and fibulas were excised. Connective tissue was removed, and the bones were sterilized with 70% ethanol. Bones were washed with BMDM media (20% HyClone FBS, 1% sodium pyruvate, 0.1% β-mercaptoethanol, 10% conditioned supernatant from 3T3 fibroblasts, in Gibco DMEM containing glucose and 100 U/ml penicillin and 100 ug/ml streptomycin) and ground using a mortar and pestle. Bone homogenate was passed through a 70 μm nylon Corning Falcon cell strainer (Thermo Fisher Scientific, 08-771-2) to remove particulates. Filtrates were centrifuged in an Eppendorf 5810R at 1,200 RPM (290 x G) for 8 min, supernatant was aspirated, and the remaining pellet was resuspended in BMDM media. Cells were then plated in non-TC-treated 15 cm petri dishes (at a ratio of 10 dishes per 2 femurs/tibias) in 30 ml BMDM media and incubated at 37° C. An additional 30 ml was added 3 d later. At 7 d the media was aspirated, and cells were incubated at 4°C with 15 ml cold PBS (Gibco, 10010-023) for 10 min. BMDMs were then scraped from the plate, collected in a 50 ml conical tube, and centrifuged at 1,200 RPM (290 x G) for 5 min. The PBS was then aspirated, and cells were resuspended in BMDM media with 30% FBS and 10% DMSO at 10^7^ cells/ml. 1 ml aliquots were stored in liquid nitrogen.

### Infections *in vitro*

To plate cells for infection, aliquots of BMDMs were thawed on ice, diluted into 9 ml of DMEM, centrifuged in an Eppendorf 5810R at 1,200 RPM (290 x G) for 5 minutes, and the pellet was resuspended in 10 ml BMDM media without antibiotics. The number of cells was counted using Trypan blue (Sigma, T8154) and a hemocytometer (Bright-Line), and 5 x 10^5^ cells were plated into 24-well plates. Approximately 16 h later, 30% prep *R. parkeri* were thawed on ice and diluted into fresh BMDM media to the desired concentration (either 10^6^ PFU/ml or 2×10^5^ PFU/ml). Media was then aspirated from the BMDMs, replaced with 500 µl media containing *R. parkeri*, and plates were spun at 300 G for 5 min in an Eppendorf 5810R. Infected cells were then incubated in a humidified CEDCO 1600 incubator set to 33°C and 5% CO_2_ for the duration of the experiment. For treatments with recombinant mouse IFN-β, IFN-β (PBL, 12405-1) was added directly to infected cells immediately after spinfection. L-BSO was obtained from Sigma (B2515).

Titers were determined via plaque assays by serially diluting the bacteria in 12-well plates containing confluent Vero cells. Plates were then spun for 5 min at 300 x G in an Eppendorf 5810R centrifuge. At 24 hpi, the media from each well was aspirated and the wells were overlaid with 2 mL/well DMEM with 5% FBS and 1.2% Avicel^®^ PH-101 (Sigma,11363). At 7 dpi, the wells were overlaid with 2 mL/well of 7% Formaldehyde (VWR,10790-708) for 30 minutes. Media was then aspirated, and the wells were stained with 0.5 mL/well of 1x Crystal Violet (VWR, 470337-534) for 15 minutes. Crystal Violet was then washed off using DI water and plaques were counted 24h later.

### Microscopy

For immunofluorescence microscopy, 2.5 x 10^5^ Vero, HMEC-1, or BMDMs were plated overnight in 24-well plates with sterile 12 mm coverslips (Thermo Fisher Scientific, 12-545-80). Infections were performed as described above. At the indicated times post-infection, coverslips were washed once with PBS and fixed in 4% paraformaldehyde (Ted Pella Inc., 18505, diluted in 1 x PBS) for 10 min at room temperature. Coverslips were then washed 3 times in PBS. Coverslips were washed once in blocking buffer (1 x PBS with 2% BSA) and permeabilized with 0.5% triton X-100 for 10 min. Coverslips were incubated with antibodies diluted in 2% BSA in PBS for 30 min at room temperature. *R. parkeri* was detected using mouse anti-*Rickettsia* 14-13 or I7 (originally from Dr. Ted Hackstadt, NIH/NIAID Rocky Mountain Laboratories). Polyubiquitin was stained the FK1 antibody (Novus). Nuclei were stained with DAPI, and actin was stained with Alexa-568 phalloidin (Life Technologies, A12380). Secondary antibodies were Alexa-405 goat anti-mouse (A31553) and Alexa-488 goat anti-rabbit (A11008). Coverslips were mounted in Prolong mounting media (Invitrogen). Samples were imaged with either the Keyence BZ-X810 Inverted Microscope with 20x or a 60x oil objective or the Airyscan 2/ GaAsP function of the Zeiss LSM 900 Confocal Multiplex with Plan-Apochromat 63x/1.4 Oil DIC M27 (FWD=0.19mm) objective or the Plan-Apochromat 20x/0.8 M27 (FWD=0.55mm) objective at the UC Irvine Microscope Imaging Core. Images were processed using FIJI^49^ and brightness and contrast adjustments were applied to entire images. Images were assembled using Adobe Illustrator. Representative images are a single optical section.

### *In vitro* assays

For LDH assays, 60 µl of supernatant from wells containing BMDMs was collected into 96-well plates. 60 µl of LDH buffer was then added. LDH buffer contained: 3 µl of “INT” solution containing 2 mg/ml tetrazolium salt (Sigma I8377) in PBS; 3 µl of “DIA” solution containing 13.5 units/ml diaphorase (Sigma, D5540), 3 mg/ml β-nicotinaminde adenine dinucleotide hydrate (Sigma, N3014), 0.03% BSA, and 1.2% sucrose; 34 µl PBS with 0.5% BSA; and 20 µl solution containing 36 mg/ml lithium lactate in 10 mM Tris HCl pH 8.5 (Sigma L2250). Supernatant from uninfected cells and from cells completely lysed with 1% triton X-100 (final concentration) were used as controls. Reactions were incubated at room temperature for 20 min prior to reading at 490 nm using an Infinite F200 Pro plate reader (Tecan). Values for uninfected cells were subtracted from the experimental values, divided by the difference of triton-lysed and uninfected cells, and multiplied by 100 to obtain percent lysis. Each experiment was performed and averaged between technical duplicates and biological triplicates.

For the IFN-I bioassay, 5 x 10^4^ 3T3 cells containing an interferon-sensitive response element (ISRE) fused to luciferase^51^ were plated per well into 96-well white-bottom plates (Greneir 655083) in DMEM containing 10% FBS, 100 U/ml penicillin and 100 µg/ml streptomycin. Media was replaced 24 h later and confluent cells were treated with 2 µl of supernatant harvested from BMDM experiments. Media was removed 4 h later and cells were lysed with 40 µl TNT lysis buffer (20 mM Tris, pH 8, 200 mM NaCl, 1% triton-100). Lysates were then injected with 40 µl firefly luciferin substrate (Biosynth) and luminescence was measured using a SpectraMax L plate reader (Molecular Devices).

### Statistical analysis

Statistical parameters and significance are reported in the Figure legends. For comparing two sets of data, a two-tailed Student’s T test was performed. For comparing multiple data sets, a one-way ANOVA with multiple comparisons with Tukey post-hoc test was used for normal distributions, and a Mann-Whitney *U* test was used for non-normal distributions. Data are determined to be statistically significant when p<0.05. All data points shown in bar graphs and in animal data are distinct samples. Asterisks denote statistical significance as: *, p<0.05; **, p<0.01; ***, p<0.001; ****, p<0.0001, compared to indicated controls. For animal experiments, bars denote medians. Error bars indicate standard deviation (SD). All other graphical representations are described in the Figure legends. Statistical analyses were performed using GraphPad PRISM.

## Data availability

*R. parkeri* strains were authenticated by whole genome sequencing and are available in the NCBI Trace and Short-Read Archive; Sequence Read Archive (SRA), accession number SRX4401164.

## Additional Information

Correspondence and requests for materials should be addressed to T.P.B.

### Author contributions

H.S., T.T.V, M.D., A.P.L, and T.P.B. performed and analyzed *in vitro* and *in vivo* experiments.

T.P.B. wrote the original draft of this manuscript. Critical reading and further edits were also provided by H.S., M.D., and A.P.L. Supervision was provided by T.P.B.

